# Distance-based exclusion method for parentage analysis using microsatellites (SSR) markers

**DOI:** 10.1101/2021.01.25.428054

**Authors:** Gil-Muñoz Francisco, Abrahamsson Sara, García-Gil M Rosario

## Abstract

Genotyping mistakes represent a challenge in parental assignment where even small errors can lead to significant amounts of unassigned siblings. Different parental assignment algorithms have been designed to approach this problem. The Exclusion method is the most applied for its reliability and biological meaning. However, the resolving power of this method is the lowest for data containing genotyping errors. We introduce a new distance-based approach which we coin as Distance-Based Exclusion (DBE). The DBE method calculates the distance between the offspring haplotype and haplotype of each of the potential fathers. The father with the lowest distance is then assigned as candidate father according to a distance ratio (α). We have tested the Exclusion and DBE methods using a real dataset of 1230 offsprings subdivided into families of 25 individuals. Each family had six potential fathers and one known mother. Compared with the Exclusion method, the DBE method is able to solve 4.7% more individuals (64.4% Exclusion vs 69.1% DBE) using the most restrictive α tested without errors. DBE method can also be used together with the Exclusion method for error calculation and to further solve unassigned individuals. Using a two-step approach, we were able to assign 98.1% of the offsprings with a total predicted error of 4.7%. Considering the results obtained, we propose the use of the DBE method in combination with the Exclusion method for parental assignment.

## 1. Introduction

Microsatellites, or simple sequence repeats (SSRs) are inexpensive markers to develop and typically highly polymorphic, which makes them preferable markers to single nucleotide polymorphisms (SNPs) in plant genetic studies when a whole-genome coverage is not required (Schlötterer 2004; Varshney et al. 2005; Guichoux et al. 2011; Putman & Carbone 2014). Because of that, some authors have proposed to implement high-throughput genotyping techniques to score SSR markers in order to exploit its advantages (Pimentel et al. 2018). Three types of error have been described inherent to the nature of the SSR technology itself (Dewoody et al. 2006). Depending on the loci amplified, *Taq* polymerase can cause ‘stutter’ bands due to slipping, making the interpretation of this alleles sometimes difficult (Jones & Avise 1997; van Oosterhout et al. 2004). This problem can result in the wrong designation of a heterozygous allele as homozygous or vice-versa (Dewoody et al. 2006). The second scoring error is described as dropout of the large-allele that results from the preferential amplification of smaller alleles in a heterozygous genotype (Dewoody et al. 2006). The third type of scoring error, so called ‘null allele’ is a lack of amplification of one allele due to a mutation at the primer site, leading to erroneous designation of a locus as homozygous (Dakin & Avise 2004; Dewoody et al. 2006). The use of PCR technique can also lead to PCR artefacts associated to the complexity of the genome itself, where more than one locus can be co amplified with the same set of primers.

A common application of SSR markers is parentage assignment (Jones et al. 2010). Assuming a complete and error-free data set the Exclusion method is a suitable method to impute the paternity of each individual from an offspring population (Chakraborty et al. 1974). Exclusion method uses the Mendelian inheritance rules, and each of the possible parents is compared with each of the offsprings. If a parent fails to share at least one allele with the offspring it becomes excluded. This method is simple and effective, but one of the disadvantages is its low tolerance to scoring errors. Despite this drawback all studies aims to use this method at least partially (Jones et al. 2010).

Several methods have been developed for parental assignment implementing different ways to sort out scoring errors of individuals not fitting under exclusion based on parsimony, maximum likelihood or Bayesian approaches (Jones et al. 2010).

This work presents a new distance-based exclusion method approach that follow the Mendelian rules of inheritance. In addition, we suggest the combined use of the distance-based method with the Exclusion method. The advantages of the distance-based method compared to normal exclusion are discussed.

## 2. Materials and Methods

### 2.1 Experimental design

Controlled crosses were performed as follows, first equal amounts of pollen from five unrelated fathers were mixed. Subsequently, the pollen mixture was combined in a 50:50 proportion with pollen from one single father. Subsequently, this pollen was used to pollinize a mother tree. The same pollen mixture was used in all families, while the single fathers were chosen to share different levels of relatedness to the mother trees described by T. J Mullin (Mullin et al. 2019).

### 2.2 DNA extraction and PCR

Needles from all potential parents and from 25 seedlings per family were sampled. DNA was extracted following the CTAB protocol (Doyle & Doyle 1987). The multiplex PCR was performed as described in Ganea et al. (Ganea et al. 2015). The primers used were spac 12.5 (Soranzo et al. 1998), ctg 4363 and ctg 1376 (Chagné et al. 2004), PtTx 3107 and PtTx 4001 (Auckland et al. 2002). The reagents used were from the Type-it Microsatellite PCR Kit from QIAGEN in a total volume of 10 μl: 2.5 μl master mix (Type-it Kit, QIAGEN), 0.2 μl Q (Type-it Kit, QIAGEN), 0.4 μl H2O (Type-it Kit, QIAGEN) and 75 μg of template DNA. Primer concentrations and dyes used were the ones described in Ganea et al. (Ganea et al. 2015). The fragment analysis was carried out using capillary electrophoresis in the Beckman Coulter CEQ8000 using the DNA Size Standard Kit-400 (Beckman Coulter) with 1.5 μl PCR product diluted in 20 μl formamide.

### 2.3 SSR scoring and parental imputation

Markers were manually scored with the CEQ 8000, Genetic Analysis System, Beckman Coulter software. Once each individual was genotyped, they were grouped into families according to their common mother together with all potential fathers (see experimental design). Each group was formed by a maximum of 25 individuals.

Parental imputation procedure consisted first in the extraction of the offspring maternal allele (*A*_*OMi*_) by selecting the offspring allele (*A*_*Oi*_) whose distance was 0 from one of the two maternal alleles (*A*_*Mj*_) for each microsatellite (*n*). In case of the existence of more than one possible allele matching the maternal genotype, both were kept as potential paternal alleles. Without genotyping errors, for each *n* microsatellite:

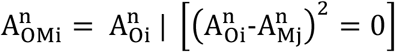

The offspring paternal alleles (*A*_*OPi*_) were the ones remaining after the extraction of the maternal alleles. The next step was to assume each potential father haploid allele (*A*_*Pj*_) set as the model alleles. The model fitness error for each potential father (*D*_Pi_) was calculated according to this formula:

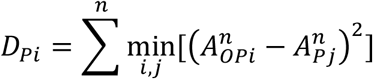

The parental models with a complete fitness (D=0) were selected as the most probable fathers. For the cases with no complete fitness (D≠0) parental imputation was conducted by taking into account the relative difference between the other candidate parents and the one with the lowest D value, a method that we coined as distance-based estimate (DBE) method.

A threshold (α) was established representing the fitness stringency for parental assignation. α=1 indicates that the father with the highest fitness value (D_P1_) is assigned irrespective of the fitness value of the next best potential father (D_P2_). Increasing values of α represents an increasing demand of fitness distance between the parent with the lowest fitness value and the next best one.

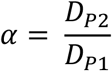

The calculated error (CE) of the DBE model for each *α* was calculated as the discrepancy between DBE and NE taken as reference parent-offspring assignation by the NE method.

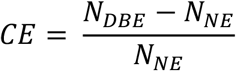

Where N_DBE_-N_NE_ is the number of mismatches between the DBE and NE methods and NE is the number of assigned individuals by the NE method. The predicted relative error (PRE) represents the predicted percentage of incorrectly assigned parents in the new data calculated by the DBE model not solved by the NE model.

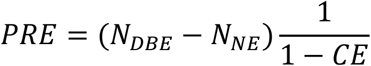

The predicted error (PE) is the predicted percentage of incorrectly assigned parents in the full model using the NE model as base and assigning the unsolved individuals using the DBE model.

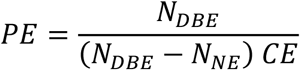

## 3. Results and Discussion

The first analysis of the results showed that comparing the obtained SSRs with the known parent genotype (mother), not all the SSRs showed an accurate allele imputation (Figure 1A). Most of the samples (87.5%) showed an accurate SSR imputation. Either by polymerase slipping or sequencing error (first type error), some errors could be identified, and this percentage proportionally decreases at greater base errorrs (Figure 1B). Above 5 bp, the error rate is higher than the previous trend, probably due to the dominancy of the other allele or failure in this allele amplification (second and third type errors).

**Figure 1.**
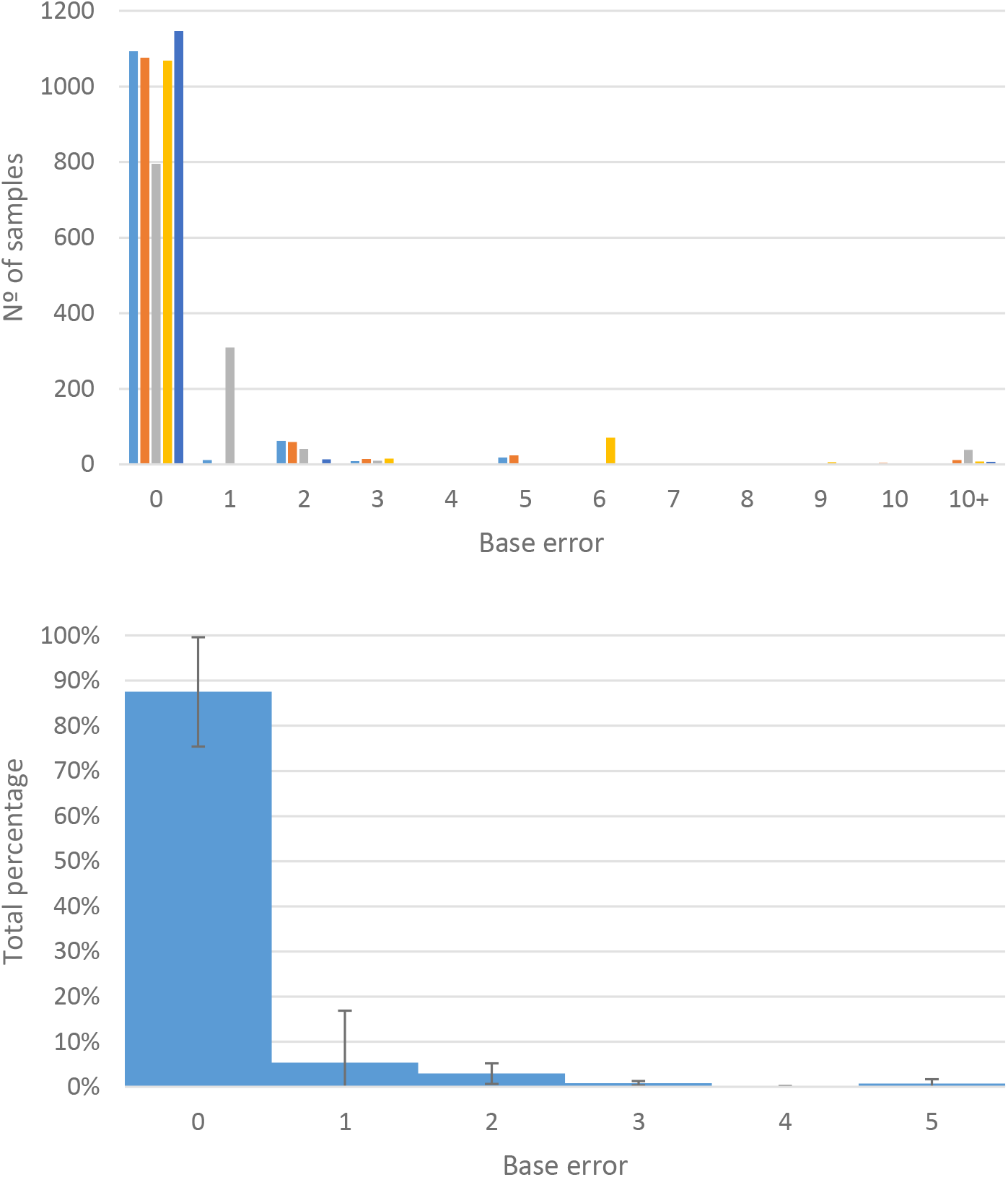

As a reference we conducted parental assignment with the exclusion (NE) method. NE method was able to solve without allele dropout 343 out of 1229 individuals. Allowing for one allele dropout in the remaining individuals resulted in the assignment of 449 additional individuals leading to a total of 792 individuals assigned. This approach solved 27.9% of the individuals without allele dropout and 64.4% when one allele dropout was allowed for the unsolved individuals (Table 1). Our results illustrate the low performance of NE method in cases where the data set contains genotyping errors, null alleles or mutations (Jones & Ardren 2003). This weakness can be palliated allowing a number of mismatches (allele dropouts) for each parent at the cost of losing reliability especially when a low number of loci is involved (Jones & Ardren 2003).

**Table 1.**

|  |  | Allele dropouts |  | Assigned<br>parents | Calculated<br>mismatches | Predicted<br>mismatches | Calculated<br>error | Predicted<br>relative<br>error | Predicted<br>error<br>NE+DBE |
| --- | --- | --- | --- | --- | --- | --- | --- | --- | --- |
|  |  | 0 | 1 |  |  |  |  |  |  |
| Normal exclusion (NE) |  | 343 | 792 | 64,4% | - | - | - | - | - |
| Distance-based<br>exclusion (DBE) | $\alpha = 1$ | 1204 | 1206 | 98,1% | 109 | 57 | 13,8% | 16,0% | 4,7% |
| | $\alpha = 1,25$ | 1050 | 1136 | 92,4% | 77 | 33 | 9,7% | 10,8% | 2,8% |
| | $\alpha = 1,5$ | 1000 | 1126 | 91,6% | 65 | 27 | 8,2% | 8,9% | 2,3% |
| | $\alpha = 2$ | 945 | 1101 | 89,6% | 55 | 21 | 6,9% | 7,5% | 1,8% |
| | $\alpha = 3$ | 875 | 1069 | 87,0% | 41 | 14 | 5,2% | 5,5% | 1,2% |
| | $\alpha = 5$ | 774 | 985 | 80,1% | 25 | 6 | 3,2% | 3,3% | 0,5% |
| | $\alpha = 10$ | 657 | 905 | 73,6% | 6 | 1 | 0,8% | 0,8% | 0,1% |
| | $\alpha = 25$ | 553 | 849 | 69,1% | 0 | 0 | 0% | 0% | 0% |

We tested the performance of the method using several values for α threshold ranging from 1 (choose the model with the closest distance) to 25. For testing the performance of the different α values we compared the obtained solutions using the DBE with the ones obtained with the NE method. The errors obtained ranged between 0% (α = 25) and 13.6% (α = 1) for our dataset (Figure 2).

**Figure 2.**
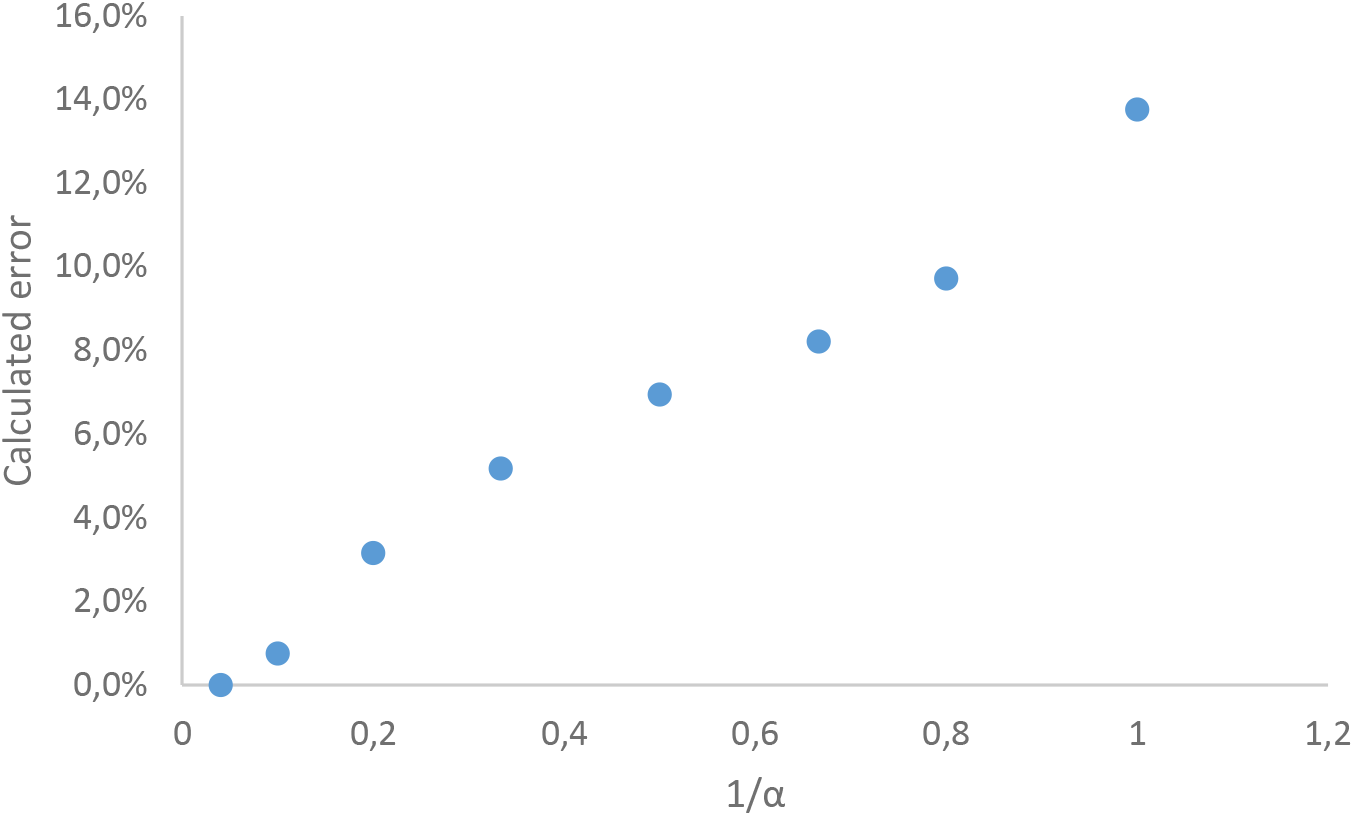

In all cases the DBE method solved a higher number of individuals compared to the NE method, even with the most restrictive α value tested (Table 1). Furthermore, the DBE method solved a higher number of individuals without allele dropout as compared to the NE method. Increasing α value also increased the proportion of individuals solved after one allele dropout (Figure 2). Solving more individuals without allele dropouts can be preferable when using a low number of alleles, as having less alleles increases the probability of assigning incorrectly an individual caused by a wrongly scored allele.

In this study, we developed a distance-based exclusion (DBE) method for parentage assignment where output distance values are generated in contrast to the NE method that is based on individuals matching. Using the NE method at first instance allows predicting the error caused by using the DBE method thus leading to a better election of the value of the cut-off parameter (α). After the application of NE, DBE method was used to solve the rest of the unsolved data (Table 1).

Both the NE and DBE methods aim for the assignment of one single parent to each individual, being the DBE method more efficient in extracting information from data sets containing scoring errors. Furthermore, the DBE method used for complement the NE method follows the Mendelian rules of inheritance and is based on the biological meaning of having one potential father in comparison to the other methods used that assign fractional parentages to each individual.

## 5. Conclusions

The DBE method, even using highly restrictive α, has been shown to have better solving capability than the NE method itself, while maintaining its biological meaning. Combined together, NE and DBE methods represent a new and powerful tool suitable for parental assignment considering that genotyping errors typically are frequent.

## Funding

This research was funded by Föreningen skogsträdsförädlingen and VINNOVA.

## Acknowledgments

We would like to acknowledge Tim Mullin, Bengt Andersson Gull and Torgny Persson for their work on the experimental design. We would also like to acknowledge the technical staff at Skogforsk and SLU that helped with the seed extraction, planting, sampling in the greenhouse and running the Beckman platform. We also acknowledge the UPSC center of Forest Biotechnology and Thomas Hiltonen for technical support on the SSR genotyping work and the Generalitat Valenciana (Spain) and European Social Fund (2014-2020) for supporting FGM PhD fellowship (ACIF/2016/115) and study visit (BEFPI/2018/056) to the department of Forest Genetics and Plant Physiology, SLU, Umeå, Sweden. We also acknowledge the contribution of Dr Laura Stefana Ganea to the microsatellite genotyping work.

## Conflicts of Interest

The authors declare no conflict of interest. The funders had no role in the design of the study; in the collection, analyses, or interpretation of data; in the writing of the manuscript, or in the decision to publish the results.

## Notes

### Competing Interest Statement

The authors have declared no competing interest.

## References

Auckland L, Bui T, Zhou Y, Shepherd M, Williams C. 2002. Conifer Microsatellite Handbook Corporate Press. College Station, TX, USA Texas A & M. University.

Björklund M. 2005. A method for adjusting allele frequencies in the case of microsatellite allele drop-out. Mol Ecol Notes.

Chagné D, Chaumeil P, Ramboer A, Collada C, Guevara A, Cervera MT, Vendramin GG, Garcia V, Frigerio JM, Echt C, et al. 2004. Cross-species transferability and mapping of genomic and cDNA SSRs in pines. Theor Appl Genet. 109:1204–1214.

Chakraborty R, Shaw M, Schull WJ. 1974. Exclusion of paternity: the current state of the art. Am J Hum Genet.

Dakin EE, Avise JC. 2004. Microsatellite null alleles in parentage analysis. Heredity (Edinb).

Dewoody J, Nason JD, Hipkins VD. 2006. Mitigating scoring errors in microsatellite data from wild populations. Mol Ecol Notes. 6:951–957. Available from: http://doi.wiley.com/10.1111/j.1471-8286.2006.01449.x

Doyle JJ, Doyle JL. 1987. A rapid DNA isolation procedure for small quantities of fresh leaf tissue. Phytochem Bull. 19:11–15.

Ganea S, Ranade SS, Hall D, Abrahamsson S, García-Gil MR. 2015. Development and transferability of two multiplexes nSSR in Scots pine (Pinus sylvestris L.). J For Res. 26:361–368.

Guichoux E, Lagache L, Wagner S, Chaumeil P, Léger P, Lepais O, Lepoittevin C, Malausa T, Revardel E, Salin F, Petit RJ. 2011. Current trends in microsatellite genotyping. Mol Ecol Resour.

Jones AG, Ardren WR. 2003. Methods of parentage analysis in natural populations. Mol Ecol.

Jones AG, Avise JC. 1997. Microsatellite analysis of maternity and the mating system in the Gulf pipefish Syngnathus scovelli, a species with male pregnancy and sex&-role reversal. Mol Ecol. 6:203–213. Available from: https://onlinelibrary.wiley.com/doi/abs/10.1046/j.1365-294X.1997.00173.x

Jones AG, Small CM, Paczolt KA, Ratterman NL. 2010. A practical guide to methods of parentage analysis. Mol Ecol Resour. 10:6–30. Available from: http://doi.wiley.com/10.1111/j.1755-0998.2009.02778.x

Mullin TJ, Persson T, Abrahamsson S, Andersson Gull B. 2019. Effects of inbreeding depression on seed production in Scots pine. Can J For Res [Internet].:cjfr-2019-0049. Available from: http://www.nrcresearchpress.com/doi/10.1139/cjfr-2019-0049

van Osterhout C, Hutchinson WF, Wills DPM, Shipley P. 2004. micro-checker: software for identifying and correcting genotyping errors in microsatellite data. Mol Ecol Notes [Internet]. 4:535–538. Available from: http://doi.wiley.com/10.1111/j.1471-8286.2004.00684.x

Pimentel JSM, Carmo AO, Rosse IC, Martins AP V., Ludwig S, Facchin S, Pereira AH, Brandão-Dias PFP, Abreu NL, Kalapothakis E. 2018. High-Throughput Sequencing Strategy for Microsatellite Genotyping Using Neotropical Fish as a Model. Front Genet [Internet]. 9. Available from: http://journal.frontiersin.org/article/10.3389/fgene.2018.00073/full

Putman AI, Carbone I. 2014. Challenges in analysis and interpretation of microsatellite data for population genetic studies. Ecol Evol.

Schlötterer C. 2004. The evolution of molecular markers - Just a matter of fashion? Nat Rev Genet.

Soranzo N, Provan J, Powell W. 1998. Characterization of microsatellite loci in Pinus sylvestris L. Mol Ecol [Internet]. 7:1260–1. Available from: http://www.ncbi.nlm.nih.gov/pubmed/9734086

Varshney RK, Graner A, Sorrells ME. 2005. Genic microsatellite markers in plants: features and applications. Trends Biotechnol [Internet]. 23:48–55. Available from: https://linkinghub.elsevier.com/retrieve/pii/S0167779904003221

